# Sleep-like state during wakefulness induced by psychedelic 5-MeO-DMT in mice

**DOI:** 10.1101/2022.12.11.519886

**Authors:** Benjamin J B Bréant, José Prius Mengual, Anna Hoerder-Suabedissen, Jasmin Patel, David M Bannerman, Trevor Sharp, Vladyslav V Vyazovskiy

**Affiliations:** Department of Physiology, Anatomy and Genetics, University of Oxford, Oxford, UK; Sleep and Circadian Neuroscience Institute, University of Oxford, UK; The Kavli Institute for Nanoscience Discovery, University of Oxford, UK; Department of Experimental Psychology, University of Oxford, Oxford, UK; Department of Pharmacology, University of Oxford, Oxford, UK

## Abstract

Psychedelics lead to profound changes in subjective experience and behaviour, which are typically conceptualised in psychological terms rather than corresponding to an altered brain state or a distinct state of arousal. Here, we performed chronic electrophysiological recordings from the cortex concomitant with pupillometry in freely moving adult male mice following an injection of a short-acting psychedelic 5-methoxy-N,N-dimethyltryptamine (5-MeO-DMT). We observed an acute induction of a dissociated state of arousal, characterised by prominent sleep-like slow waves in the cortex and marked pupil dilation in behaviourally awake, moving animals. REM sleep was markedly suppressed, similar to the effect of conventional antidepressants. We argue that the occurrence of a dissociated brain state combining features of waking and sleep may fundamentally underpin the known and hypothesised effects of psychedelics — from dream-like hallucinations to reopening of the critical period for plasticity.

## Introduction

States of arousal determine many aspects of behaviour and sensory function critical for survival. Typically, brain states are classified based on electroencephalographic (EEG) recordings, which allow their categorisation into wake, non-rapid eye movement (NREM) sleep, and rapid eye movement (REM) sleep ^1^. Wake is defined by a fast activity of low amplitude on the EEG, and the active behaviour of the animal. During NREM sleep, the animal is typically resting ^2,3^ and EEG is dominated by slow waves and spindles ^4^, while during REM sleep, sometimes referred to as “paradoxical” sleep, wake-like cortical EEG activity is accompanied with unresponsiveness and often profound muscle atonia ^1,5^. In addition to these three fundamental global states of arousal, there is a variety of mixed, hybrid or entirely distinct states where features of waking and sleep coexist. For example, sleep-like slow waves have been observed in awake animals after sleep deprivation ^6^, near lesion sites in patients with stroke ^7^, during hibernation ^8,9^, anaesthesia ^10–12^, brain injury, syncope ^13^ and coma ^14^, as well as immature states in neonate rodents or preterm babies ^15^. Even deafferented cortical slabs, organotypic neuronal cultures or brain slices can exhibit slow waves ^16^, which led to the proposal that they signify a so called “default state”, to which the brain always gravitates in the absence of stimulation or an excitatory input necessary to maintain an awake state ^17^.

The effects of drug intoxication are often described as a psychological state, such as in terms of altered or absent subjective experience. Perhaps the best-known example thereof are the so-called serotonin psychedelics, such as lysergic acid diethylamide (LSD), psilocybin, or 5-methoxy-N,N-dimethyltryptamine (5-MeO-DMT), and pharmacologically distinct compounds such as ketamine, which produce profound psychoactive effects in humans. These drugs are being investigated as novel treatments against major mental health disorders and have been shown in clinical trials to have a strong, beneficial long-term effect on treatment-resistant depressive disorders and anxiety disorders ^18–21^. One proposed neurobiological mechanism of the therapeutic action of classical psychedelics and ketamine is their demonstrated pharmacological effect on neural plasticity, including an acute change in synaptic connectivity or “reopening” of the critical period for synaptic plasticity and learning, typical for early ontogeny ^22–25^. However, the possibility that what matters in this regard is an altered state of vigilance or arousal, induced by these compounds and corresponding to a highly plastic state, has not been considered.

The motivation for this study was to address the bidirectional relationship between psychedelics and sleep, since sleep is known to be essential for brain development, neural plasticity, and mental health ^15,26,27^. We found that in mice, 5-MeO DMT acutely induced a dissociated state of arousal, in which active behaviour and pupil dilation were associated with prominent sleep-like slow waves in the cortex. We argue that this state is, on the one hand compatible with well-known and proposed subjective effects of psychedelics, including dream-like mentation and altered perception, and on the other hand is ultimately responsible for synaptic remodelling thought to be essential for their therapeutic effects.

## Results

Following the injection of 5 mg/kg i.p. 5-MeO-DMT in freely behaving mice, we observed the emergence of sleep-like slow waves on the cortical EEG (Fig. 1.a-b). While brain signals explicitly indicated sustained NREM sleep, the animals were unequivocally awake, as confirmed by direct observations and video recording (Movie S1) (Fig. 1.b). Plotting the EEG spectrograms following vehicle and 5-MeO-DMT revealed that marked effects of the compound on cortical activity lasted less than an hour in all cases (representative example: Fig. 1.c). Specifically, we observed that while the animals were awake after the injection of 5-MeO-DMT, theta-frequency activity was replaced by slow frequencies for approximately 45 min before returning to levels comparable to vehicle (Fig. 1.c; Movie S2). Further quantitative analyses determined that in the frontal derivation, the injection of 5-MeO-DMT resulted in a significant increase in EEG slow wave activity (SWA, 0.5-4Hz) and EEG spectral power in the 15–20 Hz frequency range (Fig. 1.d). In the occipital derivation, there was a significant increase in SWA and a significant decrease in theta frequency power (Fig. 1.e). As non-oscillatory scale-free activity is considered an additional reliable marker of brain state ^28,29^, we compared the spectral slope calculated for the 20 – 30 Hz frequency range between the conditions. We found that in the frontal derivation the slope was steeper after 5-MeO-DMT treatment as compared to normal wakefulness, approaching the values of NREM sleep, while the reverse was found in the occipital derivation, suggesting regional differences in the occurrence of the dissociated state (Table S1).

**Fig. 1.**
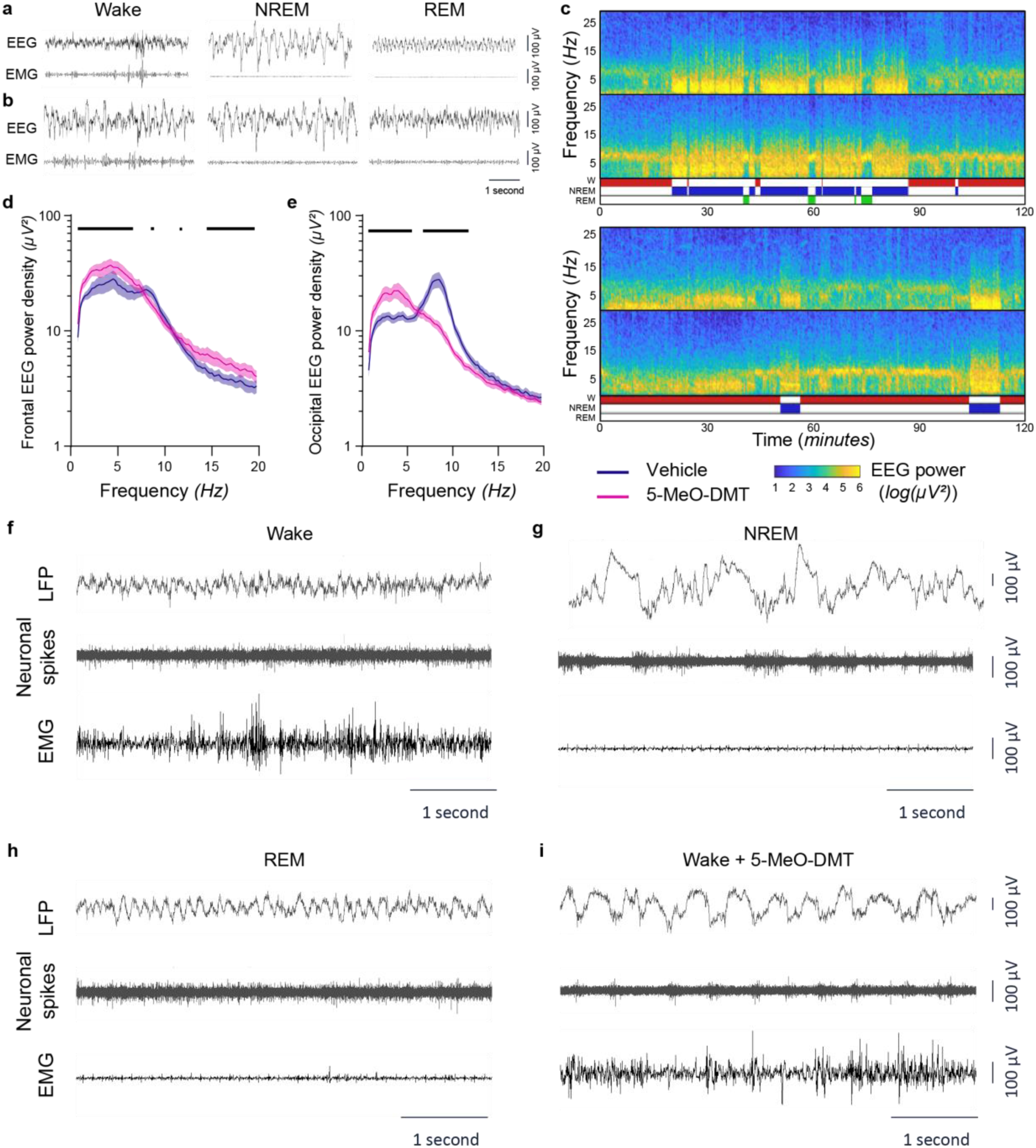
Effects of 5-MeO-DMT on brain activity. **a-b.** Representative 5-second raw EEG (top) and EMG (bottom) signal during wake, NREM and REM sleep during baseline (**a**) or within 5 minutes to 1 hour following an injection of 5mg/kg 5-MeO-DMT (**b**) showing the occurrence of slow waves during wake after an injection of 5-MeO-DMT. **c.** EEG activity shifts from theta to a broader, slower activity after an injection of 5-MeO-DMT. Representative spectrogram of the 120 minutes following an injection of vehicle (top) or 5-MeO-DMT (bottom). The top spectra of both conditions is the frontal derivation; the bottom spectra are the occipital derivations. The absence of theta rhythm and elevated slow activity is noticeable following an injection of 5-MeO-DMT. The changes in EEG activity pattern only occur within 5–45 minutes post-injection. **d-e.** An injection of 5-MeO-DMT transiently increases frontal and occipital slow wave activity and suppresses occipital theta activity. Spectral analysis of the episodes of wake occurring between 0–30 minutes after the injections in the frontal (**d**) and occipital (**e**) derivations. Differences were calculated with a ME analysis (effect of frequency*condition F_152,1060_ = 4.25, p < 0.0001, Fisher’s post hoc p < 0.05). A black horizontal line denotes a significant difference between the vehicle and 5-MeO-DMT condition for the corresponding frequency (Fisher’s LSD p < 0.05). **f-i.** An injection of 5-MeO-DMT produces sleep-like cortical activity with neuronal OFF-periods during wake behaviours. Representative LFP traces with the corresponding neuronal spiking activity and EMG activity during baseline episodes of wake (**f**), NREM (**g**) and REM sleep (**h**), or during wake in the 5–10 minutes following an injection of 5-MeO-DMT (**i**). The mean of all non-representative data is plotted with SEM.

Both NREM and REM sleep were initially suppressed after 5-MeO DMT (Fig. S1), and the animals were mostly awake during the first 1h interval after injection, but their behaviour was in many respects normal. Specifically, we never observed hyperactivity, flat body posture, catalepsy, hind limb abduction, or body tremors, and instead animals were engaged in exploratory behaviour, grooming, and nesting (Movie S1, 01:31; 03:23). It is important to note that the injection of 5-MeO-DMT did induce head twitches, behavioural changes characteristic of psychedelics (Movie S1, 01:53) (Fig. S2.a) and produced repeated exploration of the bedding immediately under the animal using their forelimbs (Movie S1). To further characterise the sensory and motivational components of wakefulness, we presented the animals with a plastic cup of sugar pellets and assessed their ability to run on a running wheel. 5-MeO-DMT increased the latency to cup interaction and pellet eating (Fig. S2.b-c; Movie S3) and shifted predominant behaviour to grooming or exploration away from wheel running that the control animals readily exhibited (Movie S4). This indicated that while the animals did not manifest obvious locomotor deficits, a degree of sensory disconnection and reduced engagement in motivated behaviours which are typical of the acute psychedelic state.

As coexistence of active behaviours and global sleep-like brain signals is unusual, we next sought to address the possible occurrence of abnormal or artefactual brain signals, unrelated to physiological slow-wave activity. To this end, we carefully inspected the local field potentials (LFP) and corresponding multi-unit activity (MUA) in the primary visual cortex (V1) recorded in a subset of animals. It is well established that slow waves during physiological sleep are accompanied by the occurrence of OFF-periods – generalised periods of synchronised neuronal silence – when the recorded populations of neurons do not generate action potentials, typically lasting 100–200 ms ^30,31^. As expected, in the vehicle condition, LFPs and MUA showed typical signatures of both wakefulness and NREM, with the latter characterised by frequent OFF-periods during LFP slow waves (Fig. 1.f-h-l, Movie S2). 5-MeO-DMT injection also resulted in the occurrence of prominent LFP slow waves during wakefulness, resembling those occurring in NREM, and invariably accompanied by neuronal OFF-periods (Fig. 1.j; Fig. S3.a, Movie S2). Furthermore, plotting the distribution of slow waves as a function of their amplitude and duration revealed a marked redistribution towards higher values in both measurements during wake after 5-MeO-DMT treatment as compared to vehicle condition (Fig. S3).

Increased SWA both in waking and sleep is typical for elevated homeostatic sleep pressure ^32^. Therefore next we addressed the effects of sleep deprivation on 5-MeO-DMT-generated slow waves. We hypothesised that if slow waves after 5-MeO-DMT were reflecting increased sleep drive, they would be enhanced when the animals are experiencing high sleep pressure. To this end, we kept a group of animals awake for four hours, followed by an injection of the same dose of 5-MeO DMT used before ^26,32–34^. As expected, we observed that NREM sleep SWA after sleep deprivation was significantly increased compared to baseline in the vehicle condition (Fig. 2.a). Interestingly, elevated sleep pressure did not result in a further increase of SWA in awake animals following 5-MeO-DMT (Fig. 2.b-c), suggesting that slow-waves induced by the drug correspond to an altered state of arousal rather than reflecting increased homeostatic sleep drive. No further difference was found in subsequent sleep episodes, besides an increase in SWA in all conditions, as typical for sleep deprivation (Fig. 2.d).

**Fig. 2.**
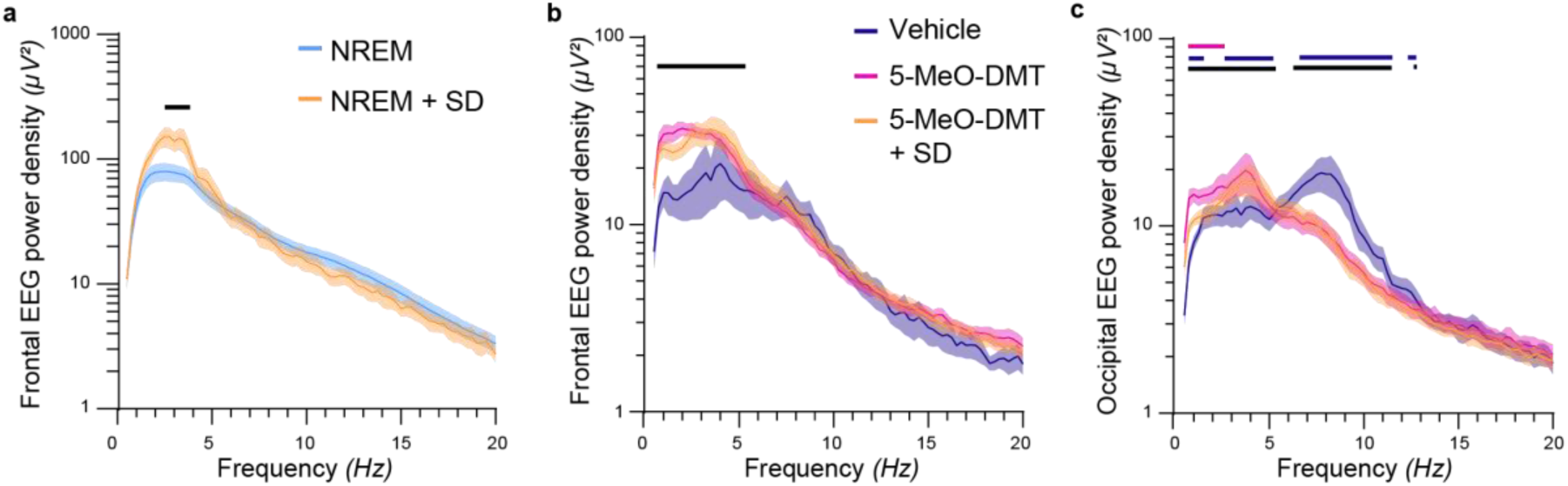
Effects of sleep homeostasis on the alterations of 5-MeO-DMT. **a.** Sleep deprivation enhances slow wave activity during the following episodes of NREM sleep. A black horizontal line denotes a significant difference between NREM and NREM+SD for the corresponding frequency (ME analysis, frequency*condition: F_234,1476_ = 6.82, p < 0.0001; Fisher’s LSD p < 0.05). **b-c.** Spectral analysis of the episodes of wake occurring between 0– 30 minutes after the injections in the frontal (**b**) and occipital (**c**) derivations. A horizontal line denotes a difference between 5-MeO-DMT and vehicle (*black*), 5-MeO-DMT+SD and vehicle (*blue*), and 5-MeO-DMT and 5-MeO-DMT+SD (*pink*) for the corresponding frequency (**b.** ME analysis, frequency*condition: F_156,1013_ = 7.40, p < 0.0001; Fisher’s LSD p < 0.05; **c.** ME analysis, effect of interaction F_156,936_ = 10.39, p < 0.0001; Fisher’s LSD p < 0.05).

The occurrence of slow waves has been linked to decreased levels of attention and alertness ^35,36^. To measure the effects of 5-MeO-DMT on global arousal, we developed a device we called an oculometer. This was based on a miniature camera previously used in birds ^37^, that was fitted to the head-stage in order to track the state of the pupil in freely behaving mice. Previous studies in mice showed that movement or exploration - behaviours typically associated with increased levels of arousal are accompanied by pupil dilation ^38–40^. In our study, unrestrained mice wearing the oculometer exhibited normal spontaneous behaviours, including exploration, grooming, nesting, and sleeping (Fig. 3.a; Movie S5), while the quality of video recordings allowed subsequent image analysis with Deep Lab Cut (Fig. 3.b) (Movie S5).

**Fig. 3.**
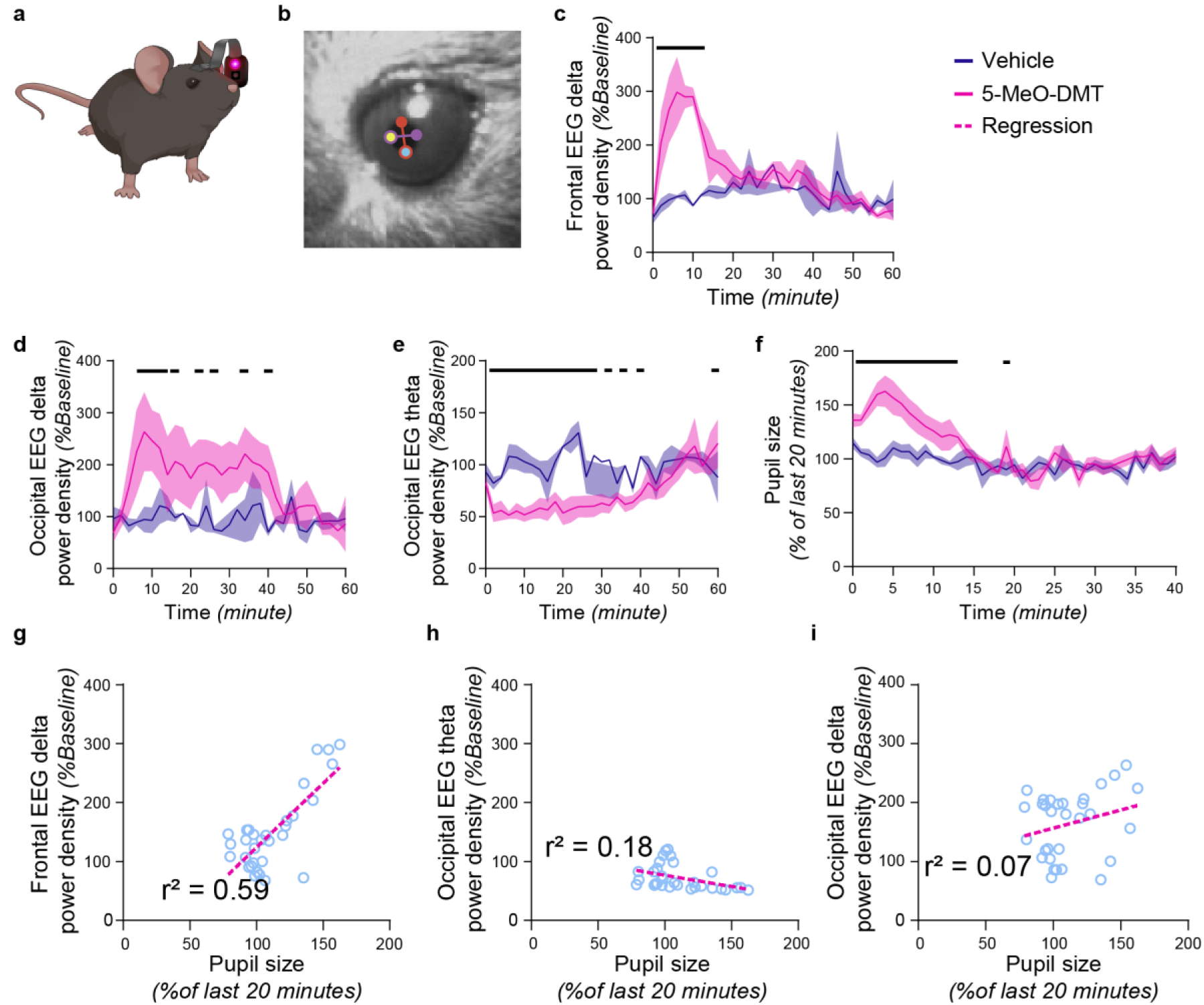
Effects of 5-MeO-DMT on cortical activity dynamics and pupil diameter. **a.** Drawing of the oculometer, a wearable device for mice capable of taking close-up video of the eyes of a freely behaving animal. **b.** Sample image from the oculometer software with the points manually scored at the north- (red), east- (purple), south- (cyan), and west- (yellow) most locations of the pupil. The North-South vector (red line) and East-West vector (purple line) used to calculate the pupil size are also represented. **c-e.** Increase in delta power and suppression of theta activity in animals wearing an oculometer are not equal, with frontal delta (**c**) increase lasting 15-20 minutes (ME analysis, effect of time point*condition; F_30,7_ = 5.19, p < 0.05; Fisher’s LSD p < 0.05), while occipital delta (**d**) and occipital theta (**e**) dynamics lasting 30–40 minutes. (**d.** ME analysis, effect of time point*condition: F_30,15_ = 2.98, p < 0.01; Fisher’s LSD p < 0.05; **e.** ME analysis, effect of time point*condition: F_30,6_ = 5.20, p < 0.001; Fisher’s LSD p < 0.05). **f**. An injection of 5-MeO-DMT is followed by an increased pupil diameter lasting 10–15 minutes. Data presented as mean + SEM (ME analysis, effect of time point*condition: F40,121 = 3.43, p < 0.0001; Fisher’s LSD p < 0.05). **g-i.** Correlation between cortical activity and pupil dilation obtained by plotting the pupil size to the EEG delta (**g-h**) and theta (**i**) activity. Pupil size is largely correlated with frontal delta activity (**g.** Pearson’s simple linear regression, r^2^ = 0.59, p < 0.0001) and negatively correlated with occipital theta activity (**h.** Pearson’s simple linear regression, r^2^ = 0.18, p < 0.05). but not occipital delta (**i.** Pearson’s simple linear regression, r^2^ = 0.07, p = 0.16).

Firstly, we replicated the effects of the injection of 5-MeO-DMT in animals wearing the oculometer, although the suppressing effects on theta-activity outlasted the effects on SWA in this cohort (Fig. S5) (Fig. 3.c-e). The injection of 5-MeO-DMT led to a marked increase (+75%) in pupil size lasting for 10–15 minutes (Fig. 3.f; Movie S6), which correlated with the increase in frontal SWA and decrease in occipital theta-power (Fig. 3.g-h). The latter was unexpected, as pupil size and elevated levels of arousal during wake behaviours have traditionally been associated with elevated hippocampal and cortical theta activity ^41,42^.

To begin addressing the origin of the changes in cortical and autonomic arousal induced by systemic administration of 5-MeO-DMT, we next performed intracortical infusions of the compound. With this experiment we tested the hypothesis that the changes in cortical neural activity were driven by local 5-HT-mediated signalling vs global changes in neuromodulatory tone. The animals were implanted with cannulae, and 500 nL of 5-MeO-DMT (4.3 mg/mL) was infused in the primary somatosensory cortex, as has been done in an earlier study with a dose of DOI that induced head twitches ^43^. Contrary to those earlier findings, we did not observe any marked behavioural or EEG alterations, although a minor increase in EEG SWA was observed after the infusion (Fig. S4). These results suggest that the prominent effects on brain state and arousal are more likely to be generated at the global level.

Unlike many serotonergic psychedelics, which preferentially target 5-HT_2A_ receptors, 5-MeO-DMT is a non-specific 5-HT receptor agonist with a 100-fold higher affinity for 5-HT_1A_ versus 5-HT_2A_ receptors ^44,45^. To investigate the role of 5-HT_1A_ receptor-mediated neurotransmission in the effects of 5-MeO DMT on brain activity, a separate group of animals was injected with 1mg/kg i.p. 5-HT_1A_ receptor antagonist WAY-100635 (WAY) 15 minutes before the injection of 5-MeO-DMT. Unexpectedly, we observed that SWA was further increased in the 5-MeO-DMT+WAY condition compared to both 5-MeO-DMT alone and vehicle conditions in both frontal (Fig. 4.a) and occipital derivations (Fig. 4.b), and this increase lasted longer than in the 5-MeO-DMT alone condition (Fig. 4.c). On the other hand, the occipital theta activity, which was strongly suppressed after 5-MeO-DMT alone, was no longer different from the vehicle condition after 5-MeO-DMT+WAY (Fig. 4.e), as was also the case for pupil diameter, whose increase was abolished by the 5-HT_1A_ antagonist (Fig. 4.f). Thus, these data dissociate 5-MeO-DMT-induced changes in frontal SWA activity and pupil diameter, with only the latter effect mediated by 5-HT_1A_ receptors (Fig. 4.g). These results suggest that different 5-HT receptor subtypes have unique and distinct contributions in different characteristics of the altered state of vigilance induced by 5-MeO DMT.

**Fig. 4.**
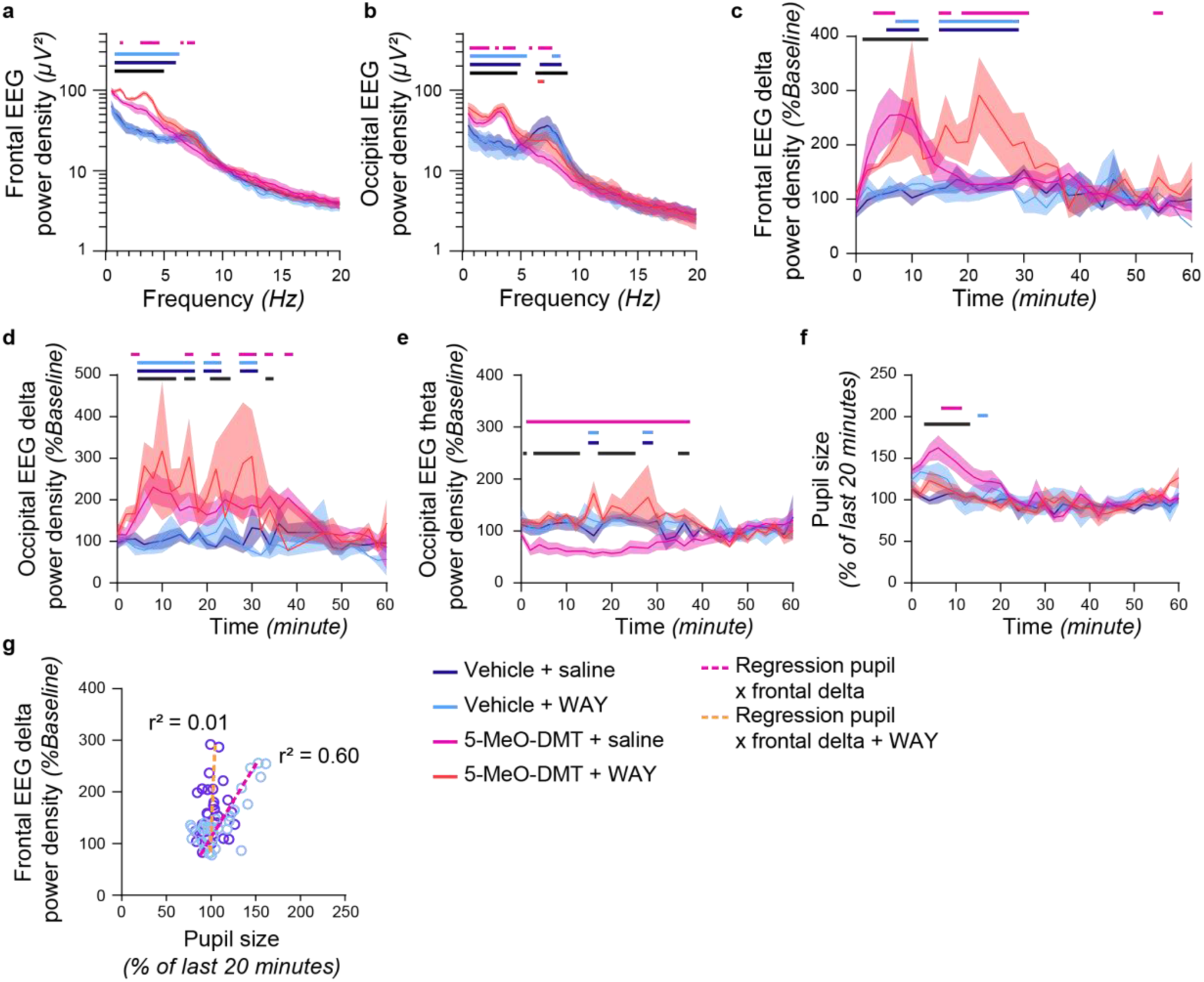
The effects of 5-MeO-DMT are mediated by 5-HT_1A_ receptors. **a.** Injection of 1 mg/kg WAY increased slow wave activity in the frontal EEG derivation when injected with 5-MeO-DMT (ME analysis, effect of frequency*condition: F_234,1009_ = 11.49, p < 0.0001, Fisher’s LSD p < 0.05). **b.** Injection of WAY did not increase occipital slow wave activity but successfully prevented the large suppression of the EEG theta activity (ME analysis, effect of frequency*condition: F_234,651_ = 13.31, p < 0.0001, Fisher’s LSD p < 0.05). **c-d.** Injection of WAY allowed a longer increase of delta activity due to 5-MeO-DMT in the frontal EEG derivation for 30–40 minutes (**c**) (ME analysis, effect of time point*condition: F_84,196_ =2.37, p < 0.0001; Fisher’s LSD p < 0.05) but had no effect regarding the time course of occipital delta activity (**d**) (ME analysis, effect of time point*condition: F_78,116_ = 2.01, p < 0.001; Fisher’s LSD p < 0.05). **e.** The suppression of the occipital theta activity occurring after the injection of 5-MeO-DMT was largely suppressed by an injection of WAY resulting in vehicle-like levels of theta activity throughout the period of interest (ME analysis, effect of time point*condition F_87,143_ = 2.92, p < 0.0001; Fisher’s LSD p > 0.05). **f.** The injection of WAY prevented the pupil mydriasis resulting from an injection of 5-MeO-DMT (ME analysis, effect of time point*condition F_120,401_ = 1.64, p < 0.001; Fisher’s LSD p > 0.05). **g**. Correlation between delta cortical activity and pupil dilation. Pupil size is largely correlated with frontal delta activity following an injection of 5-MeO-DM (Pearson’s simple linear regression, r^2^ = 0.60, p < 0.05) but not 5-MeO-DMT+WAY (Pearson’s simple linear regression, r^2^ = 0.01, p = 0.54). A red, black, pink, or dark or light blue horizontal line denotes a significant difference between: vehicle and vehicle+WAY (red, bottom); 5-MeO-DMT and vehicle (black); 5-MeO-DMT+WAY and vehicle (dark blue); 5-MeO-DMT+WAY and vehicle+WAY (light blue); and 5-MeO-DMT and 5-MeO-DMT+WAY (pink, top).

## Discussion

Here we report that 5-MeO-DMT induced a prominent change in EEG and LFP during wakefulness, as reflected in the occurrence of sleep-like slow waves associated with neuronal OFF-periods, along with marked suppression of theta-frequency activity. Despite these sleep-like patterns of brain activity, the behaviour of the animal was typical of wakefulness, and largely normal, as reflected in the occurrence of grooming, exploring, and running behaviour, as well as increased pupil diameter indicating increased arousal. The effects were short-lasting and dissipated within an hour, consistent with the kinetics of 5-MeO DMT ^45–48^. Furthermore, the 5-MeO-DMT-generated slow waves were not impacted by sleep pressure. Further investigation of potential mechanisms showed limited evidence for induction of slow waves by local application of 5-MeO DMT, and indicate that these slow waves, if anything, were enhanced in the absence of 5-HT_1A_ receptor signalling. On the other hand, the suppression of theta activity and increased pupil dilation were effectively prevented by the injection of a 5-HT_1A_ receptor antagonist.

The lack of any further increase of wake SWA when 5-MeO-DMT injection was given immediately after sleep deprivation, may suggest a ceiling effect, or can alternatively be interpreted as 5-MeO-DMT induced slow waves having distinct mechanisms, unrelated to homeostatic sleep regulation. The latter possibility is also supported by the lack of a difference during subsequent NREM sleep. We demonstrated that slow waves induced by 5-MeO-DMT are accompanied with population OFF periods, suggesting overall reduced neuronal firing, which was previously implicated in wake-dependent increase in sleep drive ^49^. Further experiments are, however, necessary to address the possibility that high wake SWA after 5-MeO-DMT provides an immediate and efficient recovery for brain networks, or simply reflects a “default”, idling cortical state where sleep pressure does not accumulate despite high levels of autonomic arousal and awake, active behaviour.

Our experiments using a 5-HT_1A_ antagonist further highlight the complexity of the mechanisms behind the effects we report in this study. We predicted that co-treatment with WAY could counteract some of the effects of 5-MeO-DMT, but initially did not expect that the main effect on cortical slow-waves would not be among those. Visual observation of the animals following co-administration of 5-MeO-DMT and WAY suggested an exacerbation of some of the 5-HT_2A_-mediated effects, such as the head twitch response ^50,51^, while wake SWA was if anything further enhanced. Since 5-HT_1A_ receptors show reciprocal interaction with 5-HT_2A_ receptors ^52^, we therefore hypothesise that the occurrence of a dissociated state with high cortical SWA is mediated primarily by 5-HT_2A_ neurotransmission. Further work is necessary to address the underlying circuit mechanisms, which could, for example, implicate cortical layer 5, known to play an important role in generation of SWA ^53^. On the other hand, our data provide important initial evidence for the functional role of slow waves, which are known to be essential for synaptic plasticity ^54^, and therefore could contribute to the proposed therapeutically-relevant effects of psychedelics ^21,25,55^.

Our data suggest that the changes in brain state and arousal are more likely to be regulated at the global rather than local cortical level. Among the wide variety of subcortical neuromodulatory systems involved in the generation of global sleep-wake states, 5-HT has a special, albeit somewhat poorly defined role. It was thought to be wake-promoting, as neural activity of 5-HT neurons in the dorsal raphe nucleus (DRN) is elevated during wake and decreases during NREM sleep, with a complete cessation of activity during REM sleep ^56–58^. However, other studies suggest a role for 5-HT in hypnogenic processes ^59,60^, and evidence is emerging that DRN 5-HT neurons can be both sleep- and wake-promoting depending on their pattern of activity ^61^. Thus, the 5-HT system is potentially rather well-placed to regulate brain state quality or intensity rather than impose a unidirectional effect of all-or-none state switching, which is consistent with the occurrence of a mixed, dissociated state induced by psychedelics.

It is well recognised that wakefulness is a highly dynamic and heterogenous state, often having features of NREM, for example during a quiet state or after sleep deprivation ^62,63^. Our current observations highlight an under-appreciated capacity of the brain to manifest hybrid states, and support the call to reconsider the relationship between SWA, vigilance states and sleep depth ^6,36,64^. Here, we show that after 5-MeO-DMT treatment, slow waves can occur concomitantly with marked pupil mydriasis, typically associated with elevated autonomic arousal ^38–40^. This suggests that local cortical mechanisms for generation and maintenance of slow-wave activity, sometimes considered a default state of brain networks ^16,65–68^, are at least partially independent from the global control of a behavioural state. We argue that the state induced by 5-MeO-DMT may not only account for its well-described subjective effects in humans, such as altered perception and ego dissolution ^69,70^, but also contribute to the proposed long-lasting therapeutic effects of psychedelics ^18,19,21,71,72^. Specifically, we propose that this property of brain networks is essential for facilitating neural plasticity, as has been shown in early ontogeny, especially in relation to so-called critical periods of development ^22,73^.

The physiological meaning of the 5-MeO-DMT-generated slow waves remains to be investigated. It has been proposed that sleep slow waves are functionally associated with structural synaptic changes, and more specifically synaptic downscaling ^54,74^. No consensus has been reached regarding the underlying cellular mechanism, but it is likely mediated by postsynaptic NMDA receptors and glycogen synthase kinase activity, coupled with rhythmic oscillations ^54^. It remains to be determined whether 5-MeO-DMT-generated slow waves actively contribute to or reflect a state permissive of synaptic downscaling ^75^, which could result in a functional decoupling and desynchronization between cortical regions ^76^. If this were the case, it could provide one highly attractive, and experimentally tractable mechanism for rapid antidepressant effects of psychedelics and other treatments associated with the induction of slow waves, such as ketamine ^10^. On the other hand, we would not rule out that the state dominated by slow waves could provide the context for synaptic plasticity occurring in both directions, such as observed during the critical periods of plasticity, especially in early development, when network slow waves are known to be prominent ^15^.

In addition to the increase of SWA, we observed that 5-MeO DMT led to a profound suppression of theta activity in the occipital derivation during active wakefulness, when it is typically present in exploring animals. Theta activity is mediated through the hippocampal formation, where rhythmical activity has been linked to waking behaviour, notably spatial and temporal processing ^41,42,77,78^. Intriguingly, human psychedelic experience has been associated with altered processing of time and space ^70^. We therefore hypothesise that suppression of theta activity observed here would be associated with a disruption of the processing of spatial and temporal information. This could include for example a dissociation between internally generated and externally generated information processing, resulting in hallucinations.

Importantly, we also observed a significant increase of REM sleep latency after 5-MeO-DMT, as was also reported after psilocin and Ayahuasca ^79–81^. Whether this effects is a direct pharmacological suppression of some of the essential defining characteristics of REM sleep, such as theta-activity, or reflects altered REM sleep regulation remains to be determined. One possibility is that the dissociated state induced by 5-MeO-DMT could replace some aspect(s) or functions(s) of REM sleep and thus reduce the homeostatic pressure to express REM sleep. REM sleep is thought to be primarily involved in emotional memory processing ^82^, which is likely relevant in psychedelic-assisted psychotherapy ^83^. We speculate that de-contextualisation of emotional memories could be facilitated during induction of a sleep-like state, associated with a reopening of critical period for neural plasticity ^22,55^.

In conclusion, we found that the injection of 5-MeO-DMT in mice induced a dissociated state of vigilance showing unequivocal features of wakefulness, such as active behaviour, high muscle tone, and dilated pupils, while brain activity was typical of NREM sleep. This paradoxical coexistence of wake and sleep characteristics mirrors that observed during paradoxical (REM) sleep and supports the notion that wake and sleep are not uniform, mutually exclusive phenomena. Furthermore, the occurrence of slow waves in awake mice and the delay in REM sleep may be key to a full understanding of the therapeutic properties of psychedelics. We speculate that the default sleep-like state emerging during 5-MeO-DMT administration has features of an immature brain state in the early ontogeny, which may manifest properties essential for heightened synaptic malleability and reopening of the critical window for plasticity ^84^, which are posited to be essential for the therapeutic effects of psychedelics.

## Methods

### Animal husbandry

Adult male C57BL/6J mice (n = 42, 9 to 15 weeks old) were kept singly housed in individual Plexiglas cages (20.3cm x 32cm x 35cm) placed inside ventilated sound-attenuated Faraday chambers (Campden Instruments, Loughborough, UK), under a 12-12 hour light-dark cycle (9 am - 9 pm). The recording room was maintained at 22 ± 1 °C and 50 ± 20 % humidity. Food and water were provided ad libitum throughout the experiment. A subset of animals was not implanted and were used to monitor head twitches (n = 4), to study the effects of injection on running wheel activity (n = 4), or to study the feeding behaviour (n = 4). All procedures were performed under a UK Home Office Project License and conformed to the Animals (Scientific Procedures) Act 1986.

### Surgeries

Procedures were performed based on established protocols for device implantation in mice ^81,85^. Prior to surgeries, mice (n = 30) were habituated to mash and jelly food and housed in individually ventilated cages. Surgeries were performed under isoflurane anaesthesia (4 % induction, 1–2 % maintenance). EEG screws were implanted above the right frontal cortex (2 mm anteroposterior, 2 mm mediolateral), right occipital cortex (anteroposterior 3.5 mm, mediolateral 2.5 mm), and left cerebellum for reference (Fig. S6.a).

In a subset of animals (n = 10), laminar probes (A1x16-3mm-100-703-Z16, NeuroNexus) were implanted in addition to EEG electrodes as above, in the primary visual cortex (−3.4 mm anteroposterior, −2 mediolateral) and referenced to the cerebellum screw.

In another subset of animals (n = 4), cannulae (C315G-005T GDE/ELE 26GA/.005“ TUNG 8637 RECEPTACLE, Bilaney Consultants Ltd) affixed with two tungsten electrodes (4.8 mm cut below pedestal; 4.7 mm and 5 mm electrode Projection) were implanted unilaterally in the primary somatosensory cortex (−1.6 mm anteroposterior; +3.3 mm mediolateral; −0.5 mm dorsoventral from brain surface, with a 30° angle). The location of the cannulae were then visually assessed by an experienced anatomist with cresyl violet coloration. EMG wires were inserted into the left and right nuchal muscles. Dental acrylic (Super Bond, Prestige Dental, Bradford, UK) was used to fix the implanted electrodes to the skull and to protect the exposed wires (Simplex Rapid, Kemdent, Swindon, UK).

A group of animals (n = 12) were also implanted frontally with a stainless-steel hexagonal nut (M2, DIN 934, RS Components) holder for the oculometer, a subset of which were also implanted with EEG and EMG electrodes (n = 6). Analgesics were administered immediately before surgery (5 mg/kg Metacam and 0.1 mg/kg Vetergesic, subcutaneous) and for at least three days following surgery (Metacam, oral). Mice were kept in individual, ventilated cages and monitored at least twice a day until baseline levels of well-being were scored for three consecutive days. They were then moved to their home Plexiglass cages for a week of habituation.

### Injections

To investigate the effects of psychedelics on vigilance states but also on behaviour and arousal, we used 5-MeO-DMT supplied by Beckley (BPL-003, Beckley Psytech, UK). 5-MeO-DMT was selected for its strong, fast-acting properties inducing dream-like experiences in humans and behavioural changes in rodents within 5 minutes of administration. Following an IP injection in mice, 5-MeO-DMT has a maximum concentration in plasma reached after 5 minutes and a terminal half-life of 12-19 minutes ^46^. On the day of the injection, 1 mg of 5-MeO-DMT was diluted into 1 mL of 0.9% sterile saline, and syringes containing either enough for an injection of 5 mg/kg 5-MeO-DMT or the vehicle (5 mL/kg sterile 0.9% saline) were prepared. The dose of 5-MeO-DMT was selected based on previous work reporting increased head twitch response at this dose ^44,48^.

After one day of baseline electrophysiological recording, the mice were subjected to a counterbalanced, crossover design whereby they received a first intraperitoneal injection at light onset (Zeitgeiber time (ZT) = 0) with either 5-MeO-DMT or vehicle and were then left undisturbed unless mentioned otherwise (Fig. S6.b). 72 hours following the first injections, the mice received an intraperitoneal injection of the substance they did not originally receive (Fig. S6.b). The majority of injected animals (n = 26) followed the above injection protocol. A similar version with the injections at a different ZT was performed for another group of animals (n = 16).

The mice implanted with cannulae (n = 4) received an intracortical injection of 5-MeO-DMT with a total volume of 500nL, at the rate of 50nL per minute. The cannulae were connected to Hamilton syringes (Microliter Syringe 700 series, 22 gauge 80621, Hamilton Company) with a connector cannula (0/SPR 40cm, Bilaney Consultants Ltd) and thin PE50 clear vinyl tubing (C232CT, Bilaney Consultants Ltd). The infusion rate of the Hamilton syringe was controlled by an automated pump (Harvard Apparatus Pump Controller, Harvard Apparatus), connected to the infuser (Nanomite Injector, Harvard Appara). As this method was novel, a concentration of 4.3 mg/mL was used based on previous work in rats using DOI and comparing the ED50 of DOI to the ED50 of 5-MeO-DMT in rats (Willins & Meltzer, 1997).

### Behavioural paradigm

To assess the effects of the compound on exploratory behaviour and feeding, animals (n = 4) received a bowl of 12 sugar pellets (Sucrose Tab/Fruit Punch 14MG, 5TUT - 1811324, TestDiet). On day 0, the bowl was introduced without prior habituation 10 minutes after dark onset (ZT12). On day 1, the animals received either a vehicle (saline) or a 5 mg/kg 5-MeO-DMT (1 mg/ml saline, Beckley Psytech) injection at dark onset (ZT = 12) at random, followed by two days of recovery, then the other injection on day 4. The bowl of sugar pellets was placed on the cage floor 10 minutes after the injections. Video recording was acquired for one hour following the injection and analysed offline.

To investigate the effects of 5-MeO-DMT on behaviour, animals (n = 4) were given access to running wheels as an assay for locomotor activity. Installation of the setup and habituation was performed based on established protocols ^86^. In a crossover design, the animals received either vehicle (saline) or a 5 mg/kg 5-MeO-DMT (1mg/ml saline, Beckley Psytech) injection at dark onset (ZT12) at random on day 1, followed by two days of recovery, then the other injection on day 4. The animals were placed in the running wheels 10 minutes following each injection and were left otherwise undisturbed to assess spontaneous activity. Video recording was acquired for one hour following the injection and analysed offline.

### Sleep Deprivation

To investigate the influence of sleep homeostasis on 5-MeO-DMT-induced changes in brain activity or to keep the animals awake during the recordings, mice (n = 8) were subjected to sleep-deprivation procedures from ZT0 to ZT4 (Fig. S6.c). At light onset, the chambers were open and nesting material removed from the home cages. Removing the nesting material introduces changes to the environment, which kept the animals awake for around 30 minutes, during which the animals explored their environment or exhibited nest-building behaviours using the bedding of the cage. Repeated nest-building behaviour was immediately followed by a manual introduction of novel objects in the cage. Novel objects included small wooden blocks, folded gloves, and various toys, which were added to the cage or replaced as required. This procedure is known to increase homeostatic sleep pressure and subsequent EEG slow-wave activity during subsequent sleep and relies on the natural exploratory behaviour of mice ^81,87,88^. The mice were continuously monitored by the experimenters, and novel objects were introduced into the environment as soon as the animal was immobile or as sleep-like slow waves became visible on the EEG.

### Pupil diameter monitoring

To monitor levels of arousal, we built a device called an oculometer consisting of an aluminium holder with a miniature camera directed at the eye (NE2D B\&W V90F2.7\_2m, NanEye, Austria) and a far-red LED (Luxeon Rebel LED 720nm LXML-PF-1, Luxeon Star Leds, Canada) (Fig. S6.d), maintained together with Sugru mouldable silicone rubber (Black, Sugru, Germany). The LED was powered through a 700 mA BuckPuck DC driver (3023-D-E-700, Luxeon Star Leds) and connected to the BioAmp Processor (RZ 2, Tucker-Davis Technologies) with an AV 9-way cable (6126423, RS Components) for activation through user input. The camera was connected to a PC via a wired board (985-NanoUSB 2.2, Mouser UK) and the video was recorded with the NanEye Viewer software (NanEye, v6.0.3, default configuration) for NanEye 2D and NanoUSB2 boards at a frame rate between 45 frames per second (fps) and 55 fps. Exposure, gain, and offset were automatically calibrated through the software to minimise loss of data due to poor lighting conditions, which can sometimes occur during some behaviours such as digging or nest building. The device was secured to the head of the mouse with a slot cheese nylon machine screw (M2 x 6 mm, DIN 84, RS Components).

To test the oculometer, the first recorded images were performed under anaesthesia (n = 3). The videos of freely moving mice (n = 12) were recorded following one 1-hour session of habituation to the device during, which all mice were fitted with their respective oculometer, and the arms of the oculometer angled to record the most optimal view of the eye. During recording days, all animals wore the device at least 10 minutes before the start of experiments. The recordings started at light onset and were followed by the injections of either vehicle or 5-MeO-DMT as described below. As the device is quite cumbersome and could impact normal behaviour of the animals, it was decided to only record 60 minutes of video per subject per session.

### Data Acquisition

Electrophysiological signals were acquired using a multichannel neurophysiology recording system (Tucker-Davis Technologies Inc., Florida, USA). Signals were acquired and processed online using the software package Synapse (Tucker-Davis Technologies Inc., Florida, USA). All signals were amplified (PZ5 NeuroDigitizer preamplifier, Tucker-Davis Technologies Inc., Florida, USA), filtered online (0.1–128 Hz) and stored with a sampling rate of 305 Hz. Signals were read into MATLAB (MathWorks) and filtered with a zero-phase Chebyshev type filter (0.5-120 Hz for EEG, 10–45 Hz for EMG), then resampled at 256 Hz.

Vigilance states were manually scored using the signals in European Data Format using SleepSign (Kissei Comtec, Nagano, Japan). The signals were divided into epochs of 4 seconds. Each epoch was scored based on visual inspections of frontal, occipital, and EMG signals (Fig. S6.e). If recorded, LFP signals were also used. Epochs with recording artefacts due to gross movements, chewing, or external electrostatic noise were assigned to the respective vigilance state but not included in the electrophysiological analysis. Wake was scored in epochs containing high EMG activity. NREM was scored in epochs containing low EMG and visual slow activity (0.5 Hz to 4 Hz) of high amplitude in the frontal or in both frontal and occipital derivations. REM was scored when the occipital signal was mostly defined by theta activity (7 Hz to 12.5 Hz) with low EMG. Epochs containing mixed frequencies of lower amplitude and high EMG between two sleep epochs (NREM–NREM or REM–NREM) were scored as movement, unless 5 or more consecutive similar epochs were scored, in which case it was classified as wake. After sleep scoring, the power spectra of the signal were computed using a fast Fourier transform routine (Hanning window) with a 0.25 Hz resolution and exported in the frequency range between 0 and 120 Hz for spectral analysis.

Fast Fourier transform signals were stored into .mat files using a custom MATLAB script containing the spectrum of each epoch and the associated vigilance state. The spectral extraction script also corrected the epoch mistakenly scored as movement when they should have been scored as wake, and *vice versa*.

For the analysis of pupil diameter, Video recordings were first converted into AVI format and analysed using DeepLabCut (Mathis et al., 2018; Nath et al., 2019). The model trained for scoring the pupil used the following parameters: resnet = 50, Shuffle 6; 5000000 trials. Four points were used to track the pupils: the northernmost, the southernmost, the easternmost and westernmost points (Fig. 3.b). Each point was marked only if it was clearly in view and not hidden by the eyelid or any light artefact. Once scored, the DeepLabCut model was trained on a randomly generated sample of images. The trained model then automatically scored all remaining videos, and only points labelled by the model with a confidence above 99% were kept. Pupil diameter was estimated as the average of the distance between the Northern and Southern point and the distance between the Eastern and Western point.

### Slow-wave detection

To investigate the incidence of local field potential (LFP), slow waves, and corresponding neuronal activity, signals from one representative animal were analysed as previously conducted ^31^. The LFP signal was first bandpass filtered between 0.5–4 Hz (stopband edge frequencies 0.2–8 Hz) with MATLAB filtfilt function exploiting a Chebyshev Type II filter design, and waves were detected as positive deflections of the filtered LFP signal between two consecutive negative deflections below the zero-crossing. Only LFP waves with a peak amplitude larger than the mean plus one standard deviation of the amplitude across all detected waves were included in subsequent analyses. Subsequently, all slow waves were aligned to their positive peak, and the corresponding average profile of neuronal spiking was computed.

### Statistical analysis

Statistical analyses were computed using Prism (GraphPad Software, Boston, Massachusetts USA, version 10.2.0 (392)). All data sets were tested for normality with the Shapiro-Wilk test, and sphericity tested with Spearman’s rank correlation test. If all assumptions were met, a parametric test was conducted; otherwise its non-parametric equivalent was selected for α = 0.05. Each figure will detail which tests were used. On figures, a significant difference is plotted as a straight black or coloured line in spectrogram analyses, as plotting the difference for each data bin would make the figure less readable. For all other figures, significance levels are indicated with black asterisks: * for p < [0.05; 0.01], ** for p <]0.01; 0.001], *** for p <]0.001; 0.0001], **** for p < 0.0001. Figures were generated using Prism and show group means with SEM for parametric data or median and interquartile ranges for non-parametric data, unless mentioned otherwise in the figure legend, such as for plotting representative raw signals of one animal.

## Acknowledgements

The authors would like to thank Laura McKillop, Christian Harding, Elise Meijer, Sian Wilcox, and Alexander Andrews for their assistance with surgery, drug preparation, equipment setup, animal husbandry, or data analysis; Gianina Ungurean and Niels Rattenborg for advice on ordering and installation of miniature cameras for monitoring pupil diameter; Stuart Peirson and Carina Pothecary on the materials and review of the oculometer; and Beckley Psytech for providing the compound for this study. The author were funded by UK Biotechnology and Biological Sciences Research Council grant (BB/M011224/1); Wellcome Trust Senior Investigator Award (106174/Z/14/Z); Wellcome Trust Strategic Award (098461/Z/12/Z); John Fell OUP Research Fund Grant (131/032); and the Medical Research Council (MR/S01134X/1).

## Author contributions

BJBB: Conceptualisation; Methodology, Investigation, Visualisation, Funding acquisition, Writing (original draft, review and editing)

JPM: Methodology, Visualisation, Writing (review and editing)

AHS: Methodology, Visualisation, Writing (review and editing)

JP: Methodology, Visualisation, Writing (review and editing)

DMB: Conceptualisation, Supervision, Writing (review and editing)

TS: Conceptualisation, Supervision, Writing (review and editing)

VVV: Conceptualisation, Investigation, Funding acquisition, Supervision, Writing (original draft, review and editing)

## Materials and Correspondence

All data and codes are available upon request.

## Supplementary information

### Supplementary figures

**Fig. S1.**
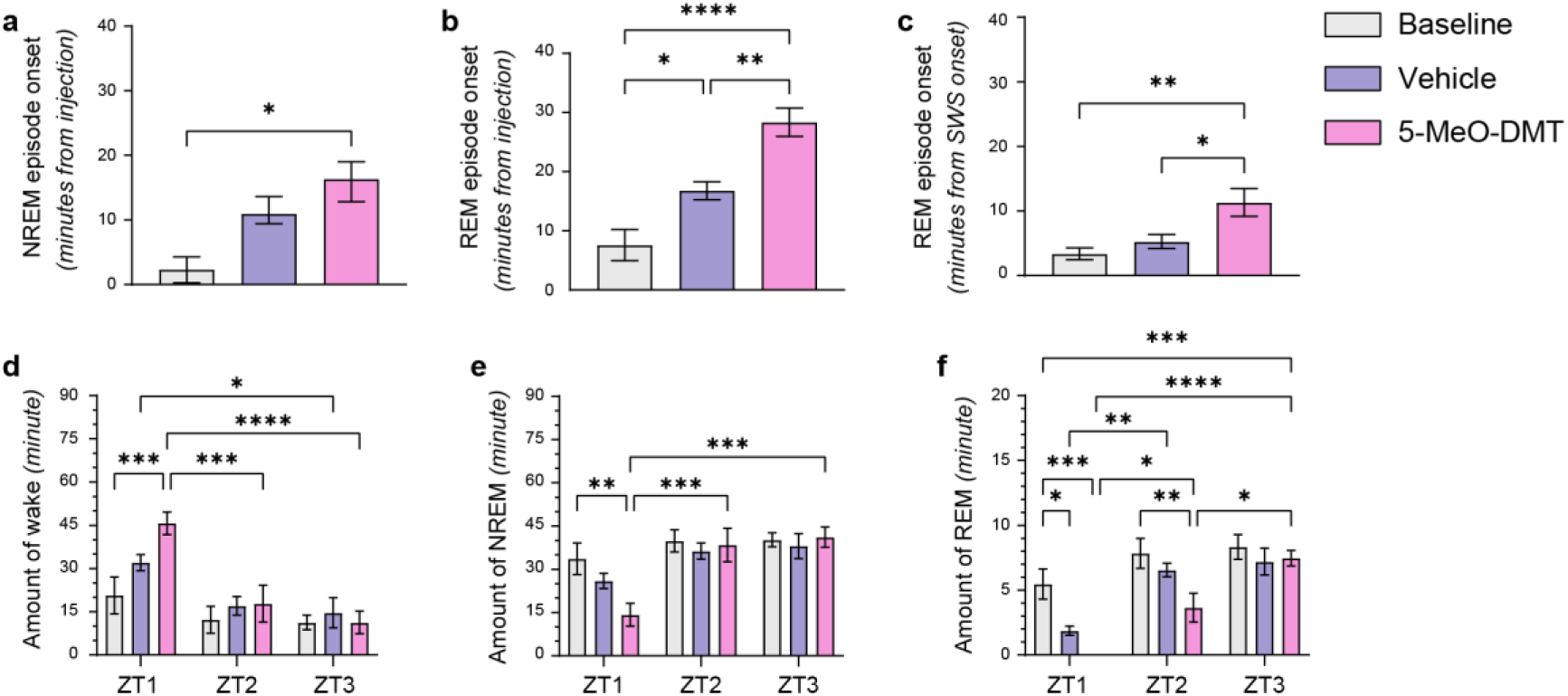
5-MeO-DMT increases NREM and REM sleep latency. **a-b.** An injection of 5-MeO-DMT increased the latency for an animal to NREM onset (**a,** median + IQRS) and REM onset (**b**) from the moment of the injection. An injection of vehicle also increased REM onset from the injection. (**a.** Friedman test χ^2^(2) = 7.75, p < 0.05; Dunn’s p < 0.05; **b.** RM-one-way ANOVA F_2,14_ = 25.22; p < 0.0001, Tukey’s p < 0.05). **c.** An injection of 5-MeO-DMT increased REM latency from NREM onset (RM-one-way ANOVA, F_2,14_ = 7.16; p < 0.01, Tukey’s p < 0.05). **d-e.** An injection of 5-MeO-DMT changes the amount of wake (**d**), NREM (**e**) and REM sleep (**f**) per hour. Wake was significantly increased in the first hour following the injection of 5-MeO-DMT, reducing the amount of NREM and REM in the same hour. The quantity of wake and NREM after 5-MeO-DMT had levels comparable to vehicle and baseline 2 hours after the injections, but the quantity of REM sleep remained significantly reduced 2 hours after 5-MeO-DMT injection. (**d.** ME analysis, effect of quantity*ZT: F_1,9_ = 210.4, p < 0.0001; **e.** ME analysis, effect of quantity*ZT: F_1,9_ = 159.1, p < 0.0001; **f.** ME analysis, effect of quantity*ZT: F_1,9_ = 193.9, p < 0.0001;). For all figures, *p<0.05; **p<0.01; ***p<0.001; ****p<0.0001. All data except a. show mean + SEM

**Fig. S2.**
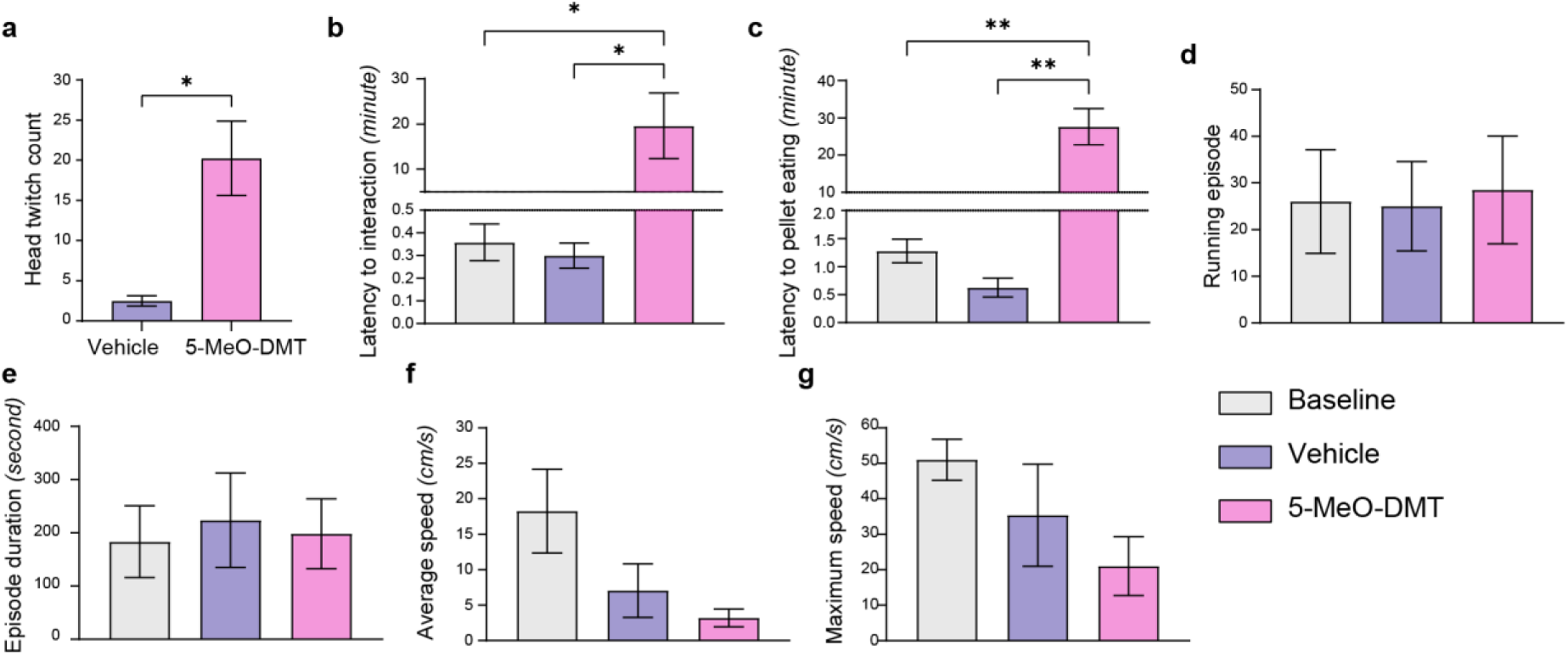
Effects of 5-MeO-DMT on wake behaviour. **a.** An injection of 5-MeO-DMT increases head twitch response from 0–60 minutes post injection (t-test, t3 = 3.96; p < 0.05). **b.** An injection of 5-MeO-DMT increased the latency for an animal to investigate a bowl of sugar pellets introduced 10 minutes after the injections. Latency is expressed from the time the bowl is deposited in the home cage and the hand of the experimenter removed from the environment (RM-one-way ANOVA, F_2,6_ = 7.07 p = 0.026; Tukey’s p < 0.05). **c.** An injection of 5-MeO-DMT increased the latency for an animal to eat a sugar pellet from the bowl (RM-one-way ANOVA, F_2,6_ = 28.95 p < 0.001; Tukey’s p < 0.01). **d-g.** Effects of an injection of 5-MeO-DMT on behaviours involving a running wheel. 5-MeO-DMT did not alter the quantity of running episodes in terms of their occurrence (**d**) or duration (**e**), with no difference in the average (**f**) or maximum (**g**) running speed (RM-one-way ANOVA, **d.** F_2,6_ = 2.854, p = 0.13; **e.** F_2,6_ = 2.16, p = 0.20; **f.** F_2,6_ = 2.81, p = 0.14; **g.** F_2,6_ = 1.81; p = 0.24). For all figures, *p<0.05; **p<0.01.

**Fig. S3.**
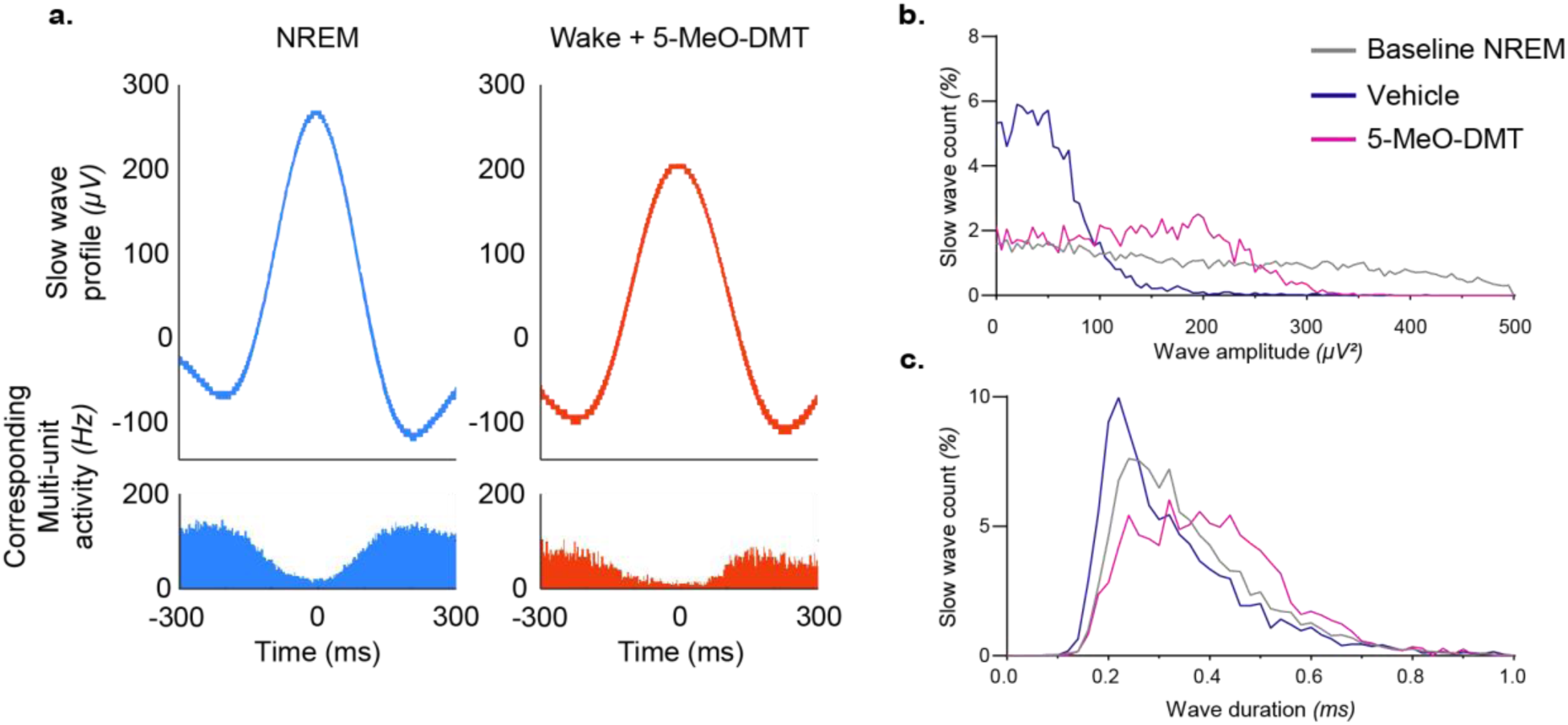
LFP slow wave analysis following an injection of 5-MeO-DMT. **a.** Representative waveform of an LFP slow wave (top) and corresponding multi-unit activity (bottom) in one representative animal during baseline NREM sleep (left) and during waking occurring 20 minutes after an injection of 5-MeO-DMT (right). **b-c.** Distribution of the amplitude (**b**) and duration (**c**) of LFP slow waves in a representative animal during NREM sleep, and during waking after the injection of vehicle or 5-MeO-DMT. Note that slow waves after 5-MeO-DMT had a higher amplitude and were longer as compared to control wakefulness.

**Fig. S4.**
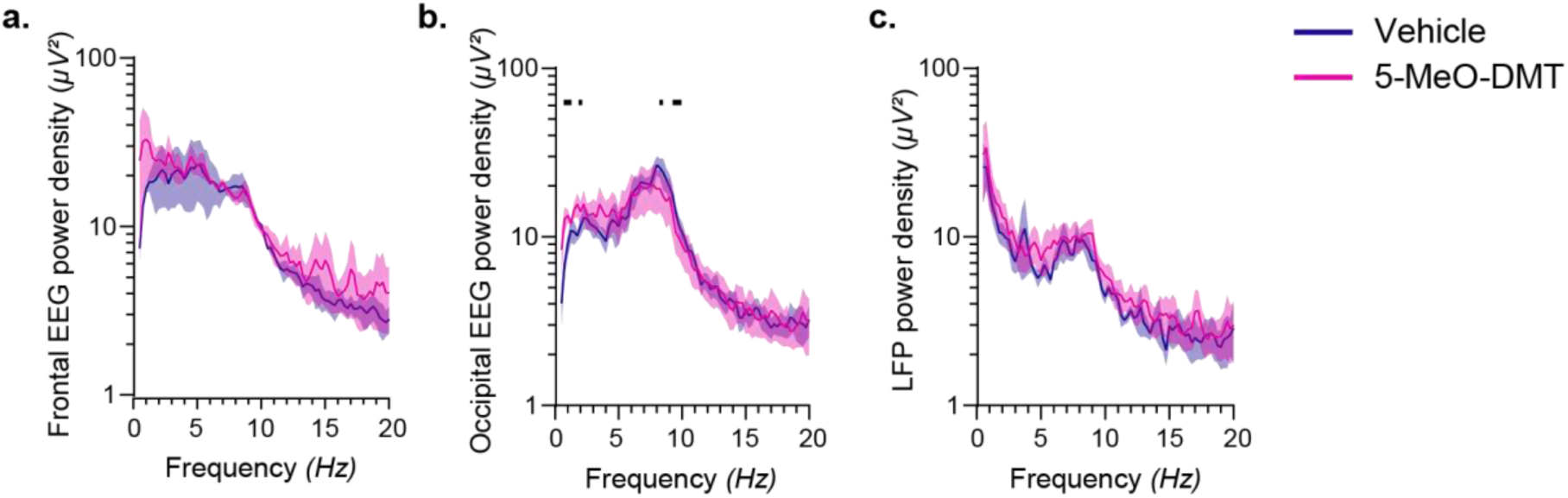
The effects of intracortical injections of 5-MeO-DMT on EEG and LFPs. **a-c**. 5-MeO-DMT intracortical injections did not induce any major significant difference in EEG frontal (**a**) and occipital (**b**) or intracortically on the LFP at the level of the injection (**c**) during the episodes of wake occurring within 30 minutes from the start of the injection. (**a.** ME analysis, effect of frequency*condition: F_156,231_ = 0.93, p = 0.68; **b.** RM two-way ANOVA effect of frequency*condition: F_156,466_ = 1.33, p < 0.05; Fisher’s LSD p < 0.05; **c.** RM two-way ANOVA effect of frequency*condition: F_156,468_ = 0.68, p = 0.99). A black horizontal line denotes a significant difference between vehicle and 5-MeO-DMT for the corresponding frequency.

**Fig. S5.**
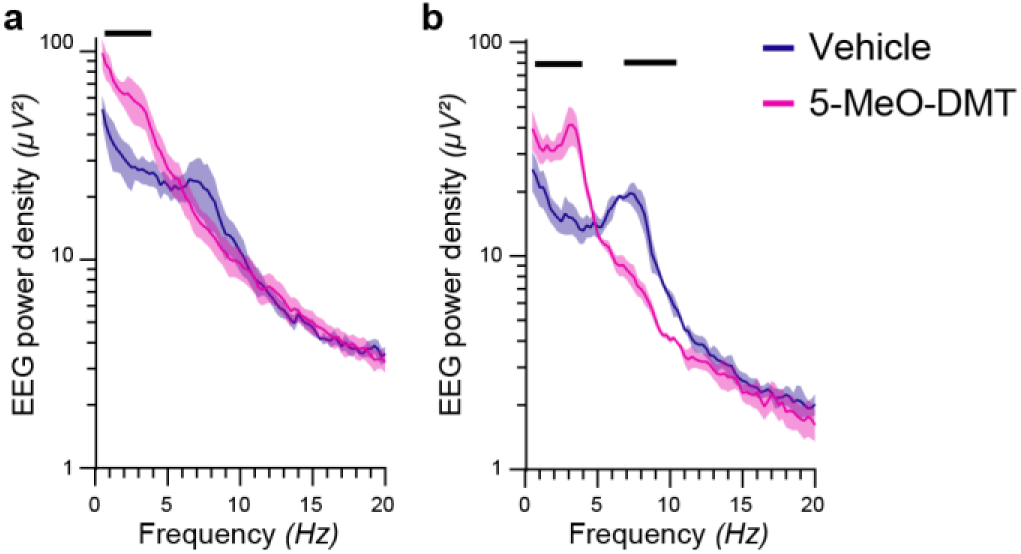
The effects of 5-MeO-DMT on the brain state of animals wearing an oculometer. **a-b**. 5-MeO-DMT injection led to a significant increase of slow wave activity during the episodes of wake occurring within 30 minutes following an injection in the frontal derivation (**a**) (ME analysis, effect of frequency*condition, F_78,75_ = 13.22, p < 0.0001; Fisher’s LSD p < 0.05) and occipital derivation (**b**), as well as a suppression of theta activity in the occipital derivation (ME analysis, effect of frequency*condition, F_78,76_ = 9.28; p < 0.0001; Fisher’s LSD p < 0.05). A black horizontal line denotes a significant difference between vehicle and 5-MeO-DMT for the corresponding frequency.

**Fig. S6.**
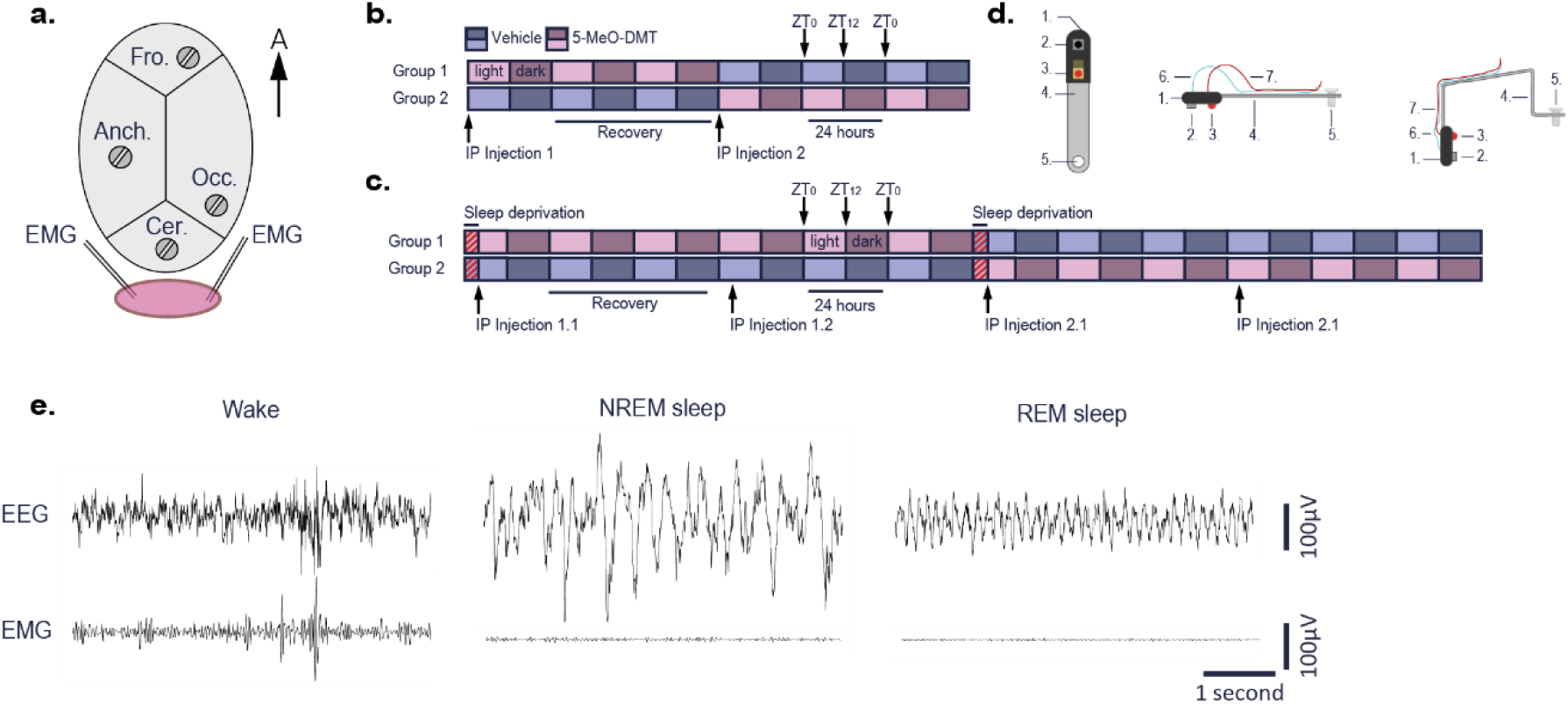
Methods. **a.** Schematic representation of a mouse skull for EEG implantation surgeries. Fro = frontal. Occ = occipital. Cer = cerebellum. Anch = anchor screw. The arrow points towards the anterior part of the animal. The EMG are implanted in the nuchal muscle. **b.** Protocol for counterbalanced, crossover injections. All animals are allocated to one of two groups. Group 1 received 5-MeO-DMT first, group 2 the vehicle. Most animals received their injection at light onset, unless mentioned otherwise. 72 hours after the first injection, the animals received a second injection with the compound they had not yet received. **c.** Sleep deprivation paradigm. The animals are sleep deprived from ZT0 to ZT4 by presenting novel objects. The injections of either 5-MeO-DMT or vehicle are made at ZT4, post-sleep deprivation. Both groups are sleep deprived at the same time. The injections 1.1 and 1.2 are the same substances and the injection 2.1 and 2.2 are the same substances, i.e.: if 1.1 was vehicle, 1.2 was vehicle and 2.1 and 2.2 were 5-MeO-DMT. **d.** Schematic of the oculometer (not to scale) facing view (left), profile view (right) unfolded (top) or folded and ready to be attached (bottom). 1. Sugru paste. 2. Camera. 3. LED. 4. Aluminium plate. 5. Screw. 6. Optic fibre. 7. Insulated copper cable. **e.** Representative EEG and EMG signals for all vigilance states. Wake is associated with EEG signals of fast frequency and low amplitude with high EMG activity. NREM is defined by large slow-waves of high amplitude in the frontal part of the brain, with low-to-absent muscle activity. In REM, the occipital EEG shows an increased theta activity due to the proximity of the electrode to the hippocampus and a flat EMG due to muscle atonia.

**Table S1.** Spectral slopes. Spectral slopes between 20 – 30 Hz for baseline wake and NREM sleep, and wake following vehicle or 5-MeO-DMT injections, with the corresponding standard error and goodness of fit. Slopes were calculated by finding the best linear fit for the spectral data between the frequencies of interest. Frontal slopes were found significantly different from one another (F_3,156_ = 97.77, p < 0.0001), as well as occipital slopes (F_3,153_ = 115.9, p < 0.0001).

| Frontal | Slope | Std. error | R<sup>2</sup> |
| --- | --- | --- | --- |
| Baseline - Wake | -0.08526 | 0.003104 | 0.9509 |
| Baseline - NREM | -0.1896 | 0.007131 | 0.9477 |
| Vehicle | -0.08832 | 0.00319 | 0.9516 |
| 5-MeO-DMT | -0.106 | 0.00527 | 0.9121 |
| Occipital |  |  |  |
| Baseline - Wake | -0.08972 | 0.002383 | 0.9732 |
| Baseline - NREM | -0.1676 | 0.005648 | 0.9292 |
| Vehicle | -0.08549 | 0.004205 | 0.9138 |
| 5-MeO-DMT | -0.06915 | 0.003181 | 0.9576 |

**Supplementary Movie 1| Waking behaviour in mice following the injection of vehicle or 5-MeO-DMT**

**Supplementary Movie 2| Representative EEG and LFP signals of a mouse in NREM sleep, REM sleep, wake with vehicle and wake with 5-MeO-DMT**

**Supplementary Movie 3| Mouse interacting with a bowl after an injection of vehicle or 5-MeO-DMT**

**Supplementary Movie 4| Mouse interacting with a running wheel after an injection of vehicle or 5-MeO-DMT**

**Supplementary Movie 5| Oculometre recording technique**

**Supplementary Movie 6| 5-MeO-DMT transiently increases pupil diameter.** The two graphs at the top are visual representation of the pupil size after vehicle (left, grey) and 5-MeO-DMT (right, red) injections against baseline (full black circle) based on the averaged pupil size over 60-minutes following an injection (bottom).

